# Tsbrowse: an interactive browser for Ancestral Recombination Graphs

**DOI:** 10.1101/2025.04.23.649987

**Authors:** Savita Karthikeyan, Ben Jeffery, Duncan Mbuli-Robertson, Jerome Kelleher

## Abstract

Ancestral Recombination Graphs (ARGs) represent the interwoven paths of genetic ancestry for a set of recombining sequences. The ability to capture the evolutionary history of samples makes ARGs valuable in a wide range of applications in population and statistical genetics. ARG-based approaches are increasingly becoming a part of genetic data analysis pipelines due to breakthroughs enabling ARG inference at biobank-scale. However, there is a lack of visualisation tools, which are crucial for validating inferences and generating hypotheses. We present `tsbrowse`, an open-source Python web-app for the interactive visualisation of the fundamental building-blocks of ARGs, i.e., nodes, edges and mutations. We demonstrate the application of `tsbrowse` to various data sources and scenarios, and highlight its key features of browsability along the genome, user interactivity, and scalability to very large sample sizes.

**Availability:** Python package: https://pypi.org/project/tsbrowse/,

Development version: https://github.com/tskit.dev/tsbrowse,

Documentation: https://tskit.dev/tsbrowse/docs/

## Introduction

Ancestral recombination graphs (ARGs) describe how a set of sample sequences relate to each other at each position along the genome in a recombining species, and are currently the subject of intense research interest (Brandt *et al*., 2024; Lewanski *et al*., 2024; Nielsen *et al*., 2024; Wong *et al*., 2024). ARGs are a fundamental object in population genetics, and although they have been of theoretical interest for decades (Hudson, 1983; Griffiths and Marjoram, 1997) it is only with recent breakthroughs in inference methods (Rasmussen *et al*., 2014; Speidel *et al*., 2019; Kelleher *et al*., 2019; Wohns *et al*., 2022; Zhang *et al*., 2023; Gunnarsson *et al*., 2024; Deng *et al*., 2024) that practical application has become possible. Varied applications have been recently proposed, such as inferring selection (Stern *et al*., 2019; Hejase *et al*., 2022) and the spatial location of genetic ancestors (Osmond and Coop, 2024; Deraje *et al*., 2024; Grundler *et al*., 2024), more powerful approaches to quantifying genetic relatedness (Fan *et al*., 2022; Zhang *et al*., 2023; Gunnarsson *et al*., 2024; Lehmann *et al*., 2025) and other methodological improvements for genome wide association studies (Nowbandegani *et al*., 2023; Link *et al*., 2023), and the development of machine learning methods using inferred ARGs as input (Hejase *et al*., 2022; Pearson and Durbin, 2023; Korfmann *et al*., 2024; Whitehouse *et al*., 2024). While these developements are exciting, the performance of these new methods depends critically on the accuracy of the inferred ARGs. Although studies benchmarking the various inference methods on simulated data have emerged (Brandt *et al*., 2022; Deng *et al*., 2024; Peng *et al*., 2024), the practicalities of applying ARG inference to real data are understudied. In particular, there is a critical lack of software infrastructure to support evaluation and quality control of inferred ARGs.

Visualisation is fundamentally important to data analysis. Many specialised tools exist to aid the visual analysis and quality control of genome assembly (Wick *et al*., 2015; Challis *et al*., 2020), read mapping (Robinson *et al*., 2011), and variant calling (Robinson *et al*., 2017; Tollefson *et al*., 2019; König *et al*., 2023), for example. At every stage of a bioinformatics pipeline, it is important to visualise results to avoid artefacts and aid understanding of the data. Genome browsers such as IGV (Robinson *et al*., 2011) and the UCSC Genome Browser (Nassar *et al*., 2023) integrate many different data modalities, and are vital infrastructure for the field.

There is currently no straightforward means of visually summarising ARGs, presenting a significant stumbling block for the nascent field of practical ARG inference. Inferred ARGs are essentially opaque, with only the most basic numerical summaries (such as numbers of nodes, mutations etc) or high-level statistics (Ralph *et al*., 2020) available. While tools for visualising the local tree topologies exist, they are difficult to interpret and do not scale well to large sample sizes. To address this gap, we present `tsbrowse`, a client-server application providing genome browser-like functionality for ARGs. It provides interactive visualisations of the information structure of ARGs, smoothly scrolling from chromosome-level views down to individual nodes, edges and mutations. Supporting very large ARGs is a particular focus for `tsbrowse`, as millions of genome sequences have been sampled for several species (Cesarani *et al*., 2022; Stark *et al*., 2024; Hunt *et al*., 2024) and ARGs of this scale are a particular focus of ongoing research (Kelleher *et al*., 2019; Zhang *et al*., 2023; Zhan *et al*., 2023; Anderson-Trocmé *et al*., 2023; Gunnarsson *et al*., 2024).

## Results

### Data model

`Tsbrowse` uses the “succinct tree sequence” encoding of ARGs (Wong *et al*., 2024). This efficient ARG encoding is implemented by the `tskit` library (Ralph *et al*., 2020) and supported by most modern ARG simulation (Kelleher *et al*., 2016, 2018; Haller *et al*., 2019; Baumdicker *et al*., 2022; Adrion *et al*., 2020; Lauterbur *et al*., 2023; Korfmann *et al*., 2023; Tsambos *et al*., 2023; Tagami *et al*., 2024), inference (Kelleher *et al*., 2019; Speidel *et al*., 2019; Wohns *et al*., 2022; Mahmoudi *et al*., 2022; Zhan *et al*., 2023; Zhang *et al*., 2023; Deng *et al*., 2025), and processing methods (Fan *et al*., 2022; Nowbandegani *et al*., 2023). In `tskit`, ARGs are encoded as a collection of tables, storing information about the nodes and edges that describe the graph topology and the sites and mutations that encode the sequence variation (Fig. 1).

**Fig. 1.**
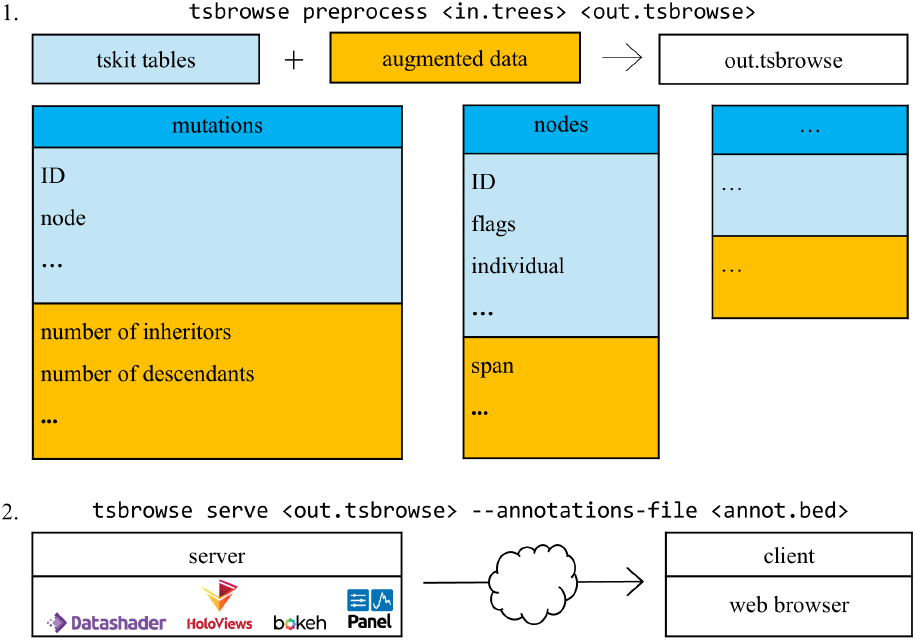
Overview of tsbrowse architecture. In `tskit`, ARGs are defined by a collection of tables. Exemplar table names are denoted in dark blue and columns that describe the property in light blue. In the pre-process step, tables are augmented with additional information computed for each property (yellow). The output from this pre-processing step is stored as a .`tsbrowse` file. Next the `serve` step renders the visualisation in a web browser by leveraging tools in the Holoviz ecosystem. Optional annotations are provided as an input file, allowing users to overlay contextual information about genes or other sequence features.

### Architecture

`Tsbrowse` is a modular client-server application written in Python, optimised for ARGs with millions of samples (Fig. 1). The application has two basic commands: `preprocess` and `serve`. The `preprocess` command takes an input ARG file and augments the `tskit` tables with additional columns, precomputing all the information required for visualisation, and storing as a compressed .`tsbrowse` file. (The .`tsbrowse` file is also a valid input for `tszip`, a general utility for compressing `tskit` files.) To visualise an ARG we then run `tsbrowse` `serve`, which by default will open a browser window on the local machine, but also supports running the server on a remote machine across the network. This client-server architecture has some important advantages over a monolithic single-machine approach. Most importantly, the ARG being visualised remains in-situ and does not need to be downloaded from the server.

The goal of `tsbrowse` is to provide interpretable and interactive visual summaries of ARGs containing millions of nodes, edges and mutations. Mutations, for example, have a clear interpretation when plotted on genome coordinate vs time axes, but at this scale the data density is far too high to simply plot each mutation as a point. We overcome this problem by using Datashader and the wider Holoviz ecosystem (Holoviz developers, 2024), which efficiently rasterises large datasets at the requested resolution on the server, sends the image to the web-browser for display, and dynamically updates as the user interactively navigates the ARG. This approach allows us to summarise very large ARGs interactively; Fig S4, for example, shows a screenshot of `tsbrowse` summarising the 1.9 million mutations in a SARS-CoV-2 ARG with around 2.7 million nodes and edges.

#### User Interface

The user interface is presented as a dashboard, with pages to describe various aspects of the ARG. Figure 2 demonstrates the user interface using the Edges page as an example. The plot on the left is an overview of the 33,929 edges in a simulated ARG with a strong selective sweep (see Sec. 4.1 for details). Each edge is depicted as a horizontal line connecting the genomic coordinates of the parent and child nodes on the x-axis, and the y-axis shows the time of either the parent or the child, as chosen by the user. The user can interact with the main “genome browser” window using a set of controls on the top-right corner (provided by Bokeh), allowing the user to pan and zoom as required. The histograms to the right then summarise the edges as depicted in the browser window. A similar browser interface is provided for nodes (e.g. Fig S3) and mutations (e.g. Fig S4). `Tsbrowse` also provides an interactive table viewer with flexible search and sorting utilities, which is a valuable debugging utility for developers.

**Fig. 2.**
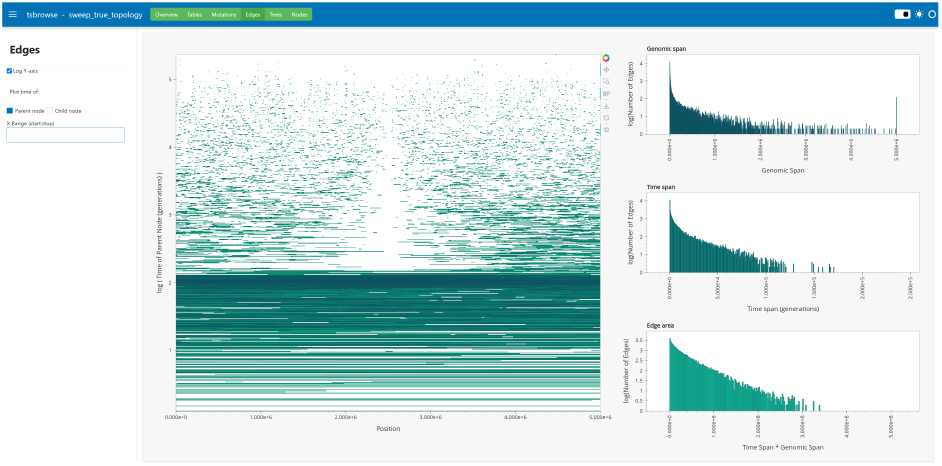
Screenshot of the Edges tab in `tsbrowse`. Visualisation of the edges in a simulated ARG with a strong selective sweep in the middle of the genome (see Supplement for details). Edges are shown in the main browser pane on the left, with additional histograms summarising the edges on the right. The effect of the sweep can be seen by the lack of edges crossing the centre of the simulated chromosome due to an excess of recent coalescent events. Moving away from the focal site, the oldest edges are the first to rejoin the ARG, followed by more recent edges, resulting in a wedge-like pattern of missing edges in the centre.

#### Applications

The purpose of `tsbrowse` is to provide an interactive view of the `tskit` ARG data model to guide intuition, improve inference quality control and facilitate debugging. An example of how we can deepen our understanding using the genome-browser-like perspective provided by `tsbrowse` is given in the simulated ARG of Fig 2, where the gap in the edges view corresponds to the characteristic dip in diversity of a selective sweep. Comparing this ground truth to the ARGs inferred by four different inference methods in Fig S1, we can see that there are substantial qualitative differences between the results. These differences in ancestral haplotypes illustrated by the edges view are unlikely to be captured by the tree-by-tree distance metrics usually used to evaluate inference methods (e.g. Kelleher *et al*., 2019; Zhang *et al*., 2023), providing further motivation for new and improved ways to compare simulated and inferred ARGs (Fritze *et al*., 2024).

Interest in ARG-based methods is burgeoning, but the methods are new and practical guidance on applying inferences to real data is lacking. Data filtering is essential, and the effects of the choices that must be made along any bioinformatics pipeline on the final ARG are hard to predict and quantify. `Tsbrowse` was primarily developed as a way to quickly visualise the effects of such filtering choices on ARGs inferred by `tsinfer`, and it is now an indispensable element of the inference pipeline. Fig S2 shows a region of the 1000G data with gaps in site density that are spanned by exceptionally long edges, which are likely to bias downstream statistics. These interactive visualisations have also helped diagnose issues with the `tsinfer` inference algorithm. Fig S3 shows the genomic spans of ancestral nodes in an ARG inferred with `tsinfer`, demonstrating a clear excess of long haplotypes in the very ancient past. These insights have helped guide development and may lead to significant improvements in performance.

## Discussion

Visualisation of tree topology is a central task in phylogenetics, and although numerous tools exist (e.g. Huson *et al*., 2007; Vaughan, 2017), the methods can typically only handle a few hundred nodes. Visualisation of large-scale tree topologies with millions of nodes requires much more sophisticated approaches to capture topological features at different scales, and is an active research area (Wong and Rosindell, 2022; Kramer *et al*., 2023). Adapting such methods, and integrating into `tsbrowse` to provide a local tree viewer that operates at the million-node scale is an important direction for future work. An interactive viewer for the entire ARG topology, capturing semantic properties of the graph at a range of scales, is an even more ambitious goal, and would be a major asset for the field.

## Supplementary Methods

### Simulation of truth dataset (Fig. 2)

Ancestral histories of 300 samples were simulated with `SweepGenicSelection` function in `msprime` (`version 1.3.3`). A combination of models was used: in the recent past, a selective sweep was simulated with a beneficial allele situated in the middle of a 5 Mb sequence. The frequency of the allele in the population was set at 0.0001 at the beginning of the sweep. The allele fixed in the population at a frequency of 0.9999. The strength of selection was set using the selection coefficient, *s = 0*.*25*. A time increment, *dt = 1e-6* was used to step through the sweep. Mutations were added to the ARG at a rate of 1e-8 per base pair per generation. A recombination rate of 1e-8 per base pair per generation. For simulating history before occurrence of the sweep, a standard coalescent model (Hudson’s algorithm (Hudson, 1983)) was used until coalescence was achieved.

### Inference of selective sweep dataset (Suppl. Fig. S1)

The following software were used to infer ARGs from the truth dataset described above: `tsinfer` version 0.3.3 (Kelleher *et al*., 2019), tsdate version 0.2.1, Relate version 1.2.2 (Speidel *et al*., 2019), ARG-needle version 1.0.3 (Zhang *et al*., 2023), SINGER version 0.1.8-beta (Deng *et al*., 2024). For all methods, the following parameters of inference were used: *recombination rate = 1e-8, mutation rate = 1e-8, effective population size = 10,000*. Default values were used for all other inference parameters.

### Inference of 1000 Genomes dataset (Suppl. Fig. S2, Suppl. Fig. S3)

The 1000 Genomes dataset was downloaded from https://ftp.1000genomes.ebi.ac.uk/vol1/ftp/data_collections/1000G_2504_high_coverage/working/20220422_3202_phased_SNV_INDEL_SV/. The ancestral fasta sequence for chromosome 17 (GRCh38) was downloaded from the Ensembl database. Inference was performed with a Snakemake pipeline (https://github.com/benjeffery/tsinfer-snakemake/) using `tsinfer` version 0.3.3 for the long arm of chromosome 17 after filtering out duplicate variant positions, variants with missing or low quality ancestral allele, singletons, n-1-tons and n-2-tons. Only bi-allelic SNPs were included for inference. For Suppl. Fig. S2, tsdate version 0.2.1 was used to estimate the age of ancestral nodes with *mutation rate = 1*.*29e-8*, setting all other parameters to default values.

### Inference of SARS-CoV-2 ARGs

The ARG shown in Fig S4 was inferred with sc2ts (Zhang *et al*., 2023) using the Viridian dataset (Hunt *et al*., 2024). It consists of 2,482,157 samples 2,689,054 nodes, 2,689,982 edges and 1,923,169 mutations. Running `tsbrowse` `preprocess` on the input `tszip` file (113M) required 2m19s time elapsed (15m15s CPU time) on an Intel Core(TM) i7-9700 CPU. The resulting .`tsbrowse` file size was 130M.

Code to recreate datasets and infer ARGs are available in an accompanying repository: https://github.com/savitakartik/tsbrowse-paper.

## Supplementary Figures

**Fig. S1.**
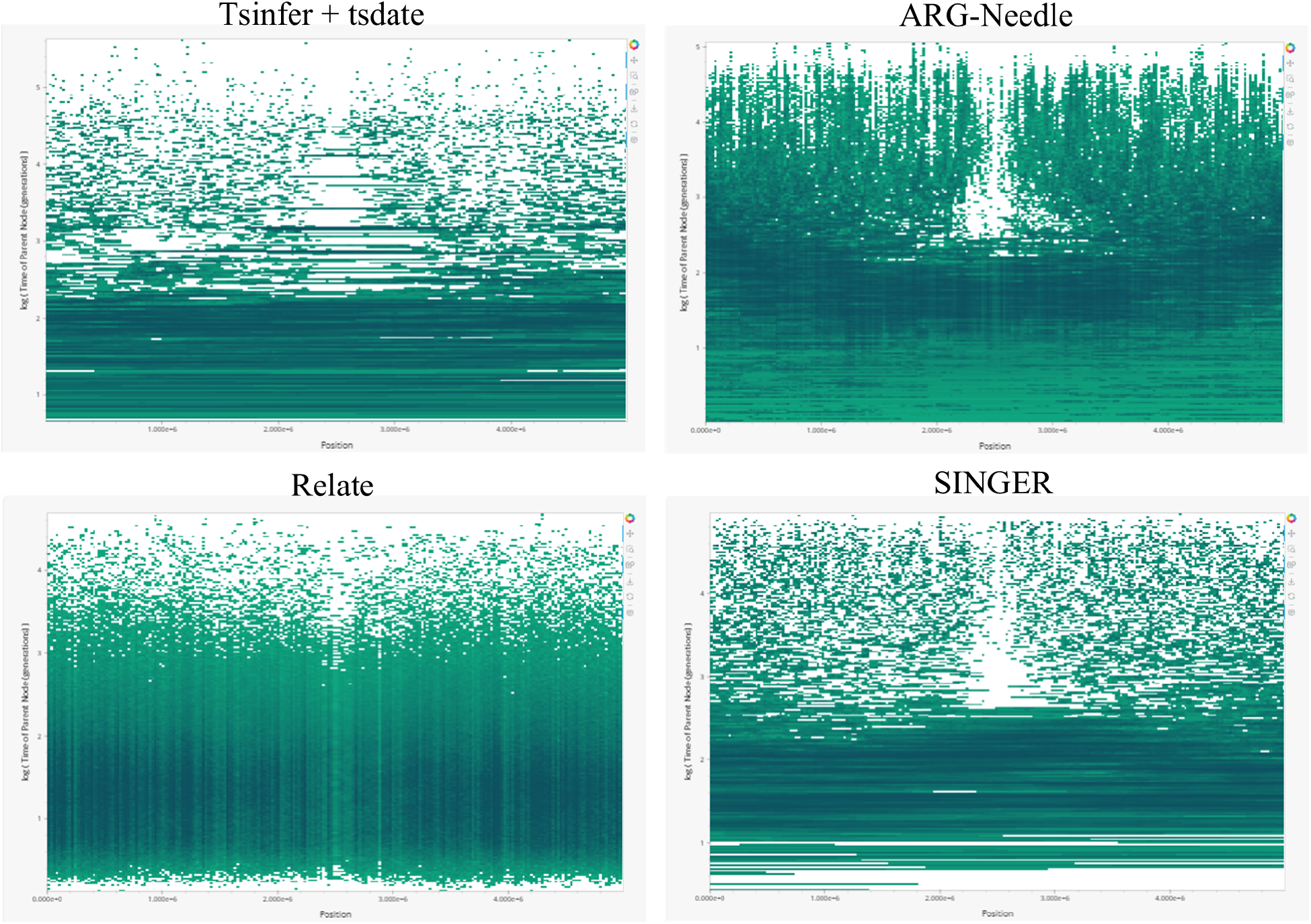
tsbrowse applied to inference methods. A screenshot of `tsbrowse`’s Edges view for `tsinfer`+tsdate, ARG-Needle, Relate and SINGER inferences of the truth dataset simulated under a selective sweep model (in Figure 2 of the main text). For SINGER, an examplar ARG from the set of output posterior ARG samples is shown. The X coordinate represents genomic position, each horizontal segment on the plot shows the genomic coordinates that the edge spans, and Y coordinate shows time of either the parent or child node in the edge.

**Fig. S2.**
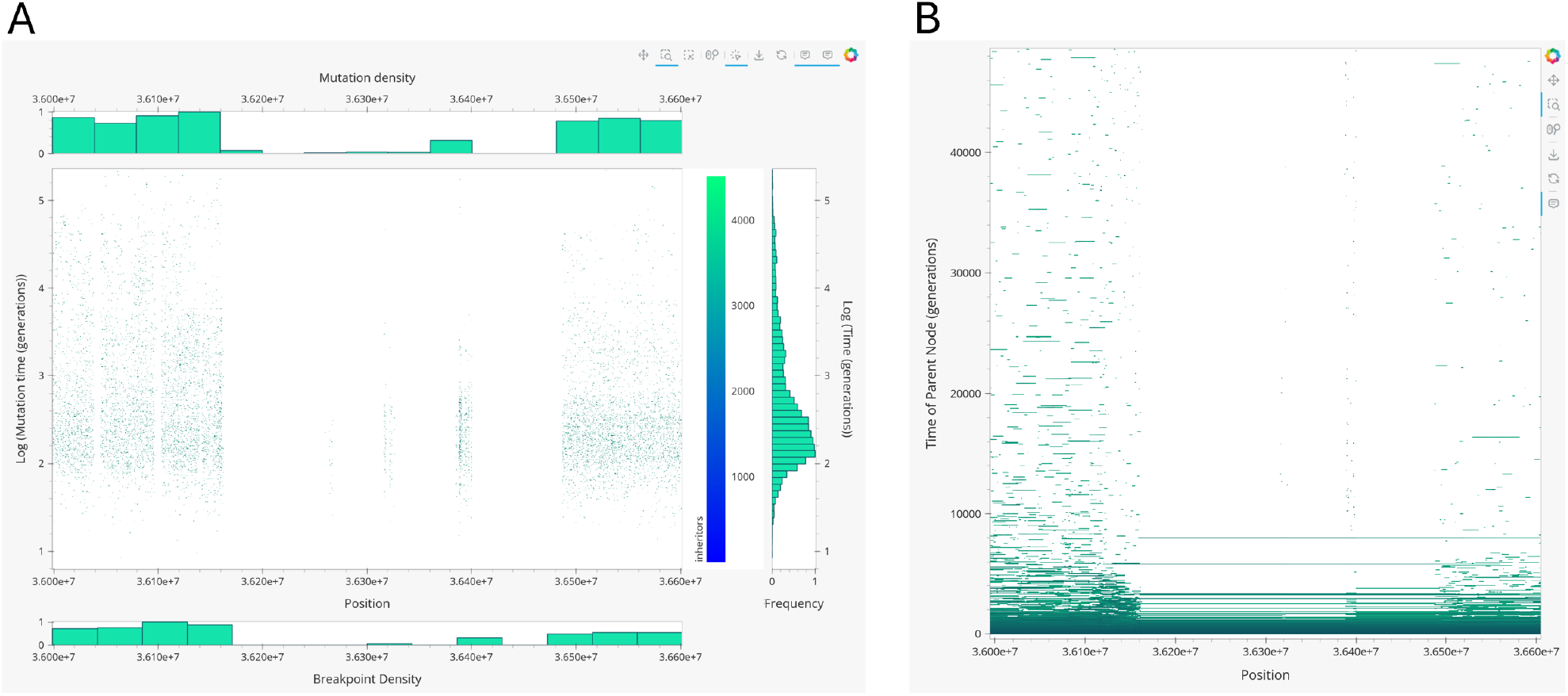
Identifying ARG inference problems with tsbrowse. Screenshots of `tsbrowse`’s Mutations view (A) and Edges view (B) for a 600 kB region of chromosome 17 inferred from 3,202 participants from the 1000 Genomes Whole Genome Sequencing dataset (1000G) (Consortium, 2015; Byrska-Bishop *et al*., 2022). The poor performance of `tsinfer` in this variant-poor region is evidenced by the long edges spanning gaps in mutation density.

**Fig. S3.**
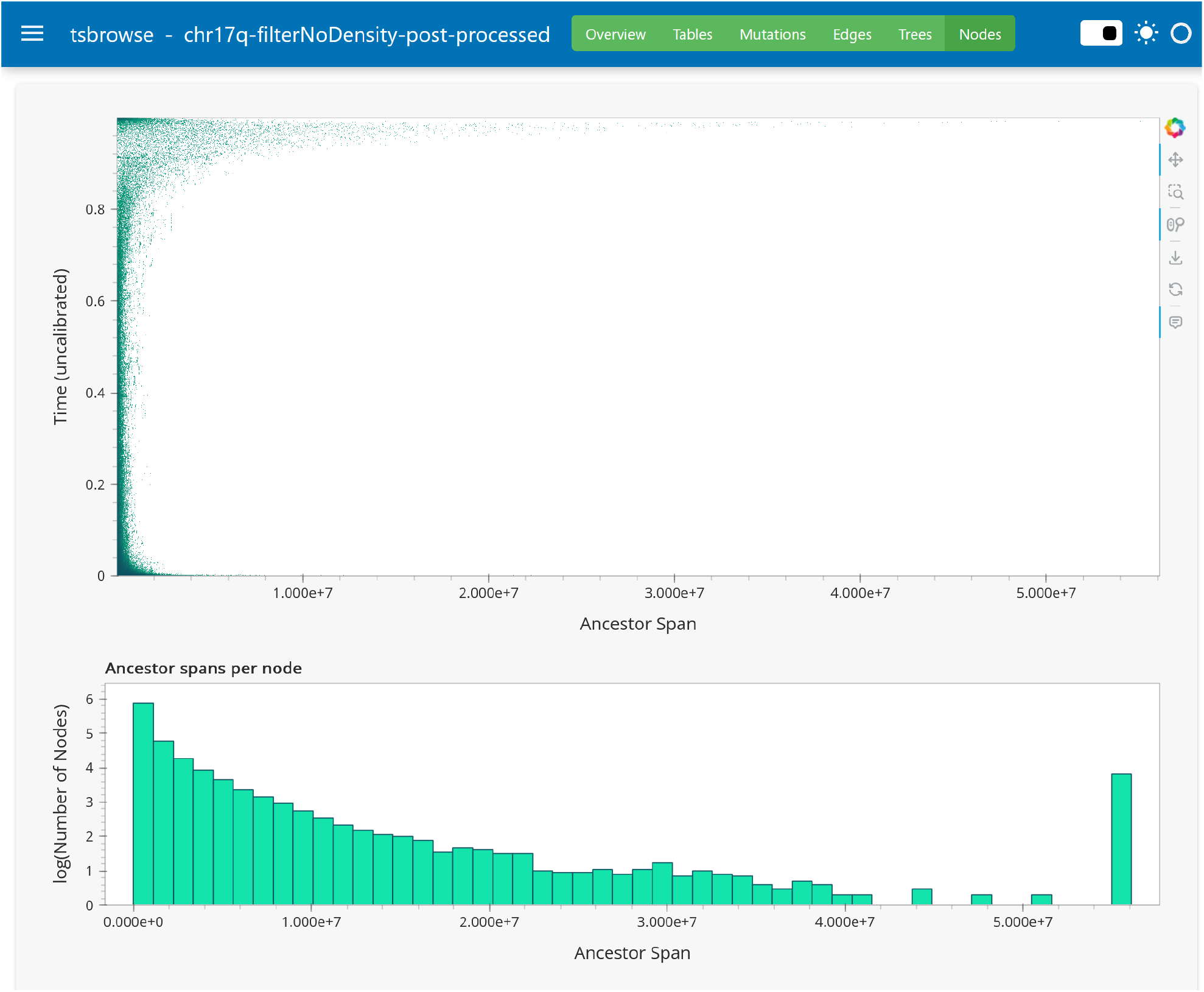
A view of the Nodes page for a 1000 Genomes inference. A screenshot of `tsbrowse`’s Nodes view for an inference of the long arm of chromosome 17 from the 1000 Genomes whole-genome sequencing dataset. At the top is a plot of node spans over time. The length of sequence that the nodes span is shown on the X axis, and the time of nodes is shown on the Y axis. The histogram at the bottom shows the distribution of node spans.

**Fig. S4.**
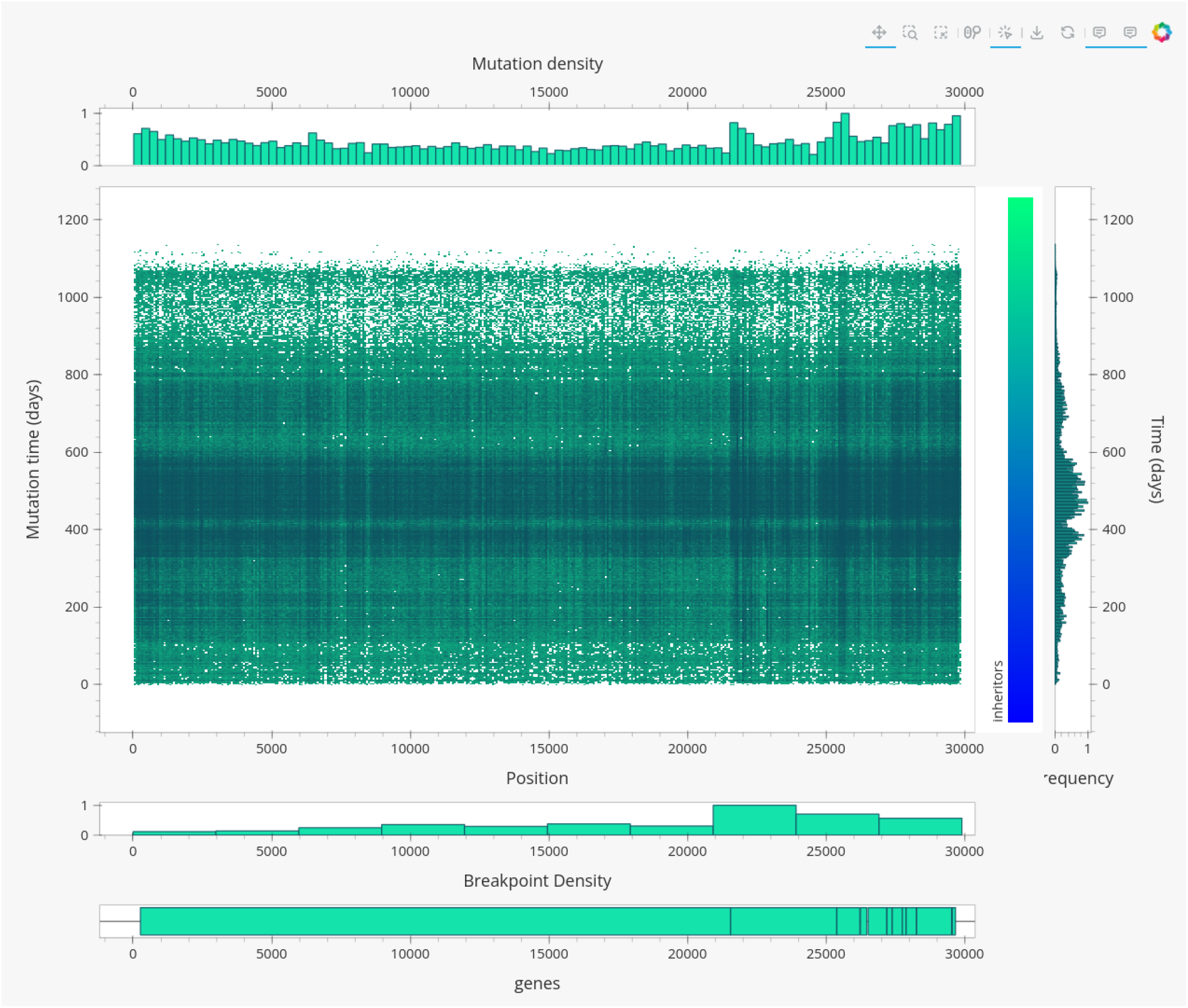
tsbrowse applied to SARS-CoV-2 ARGs. A screenshot of `tsbrowse`’s depiction of 1,923,169 mutations in an ARG inferred by sc2ts; see text for details. Also shown are the gene annotations along the X-axis.

